# Spatial Coordination between Leaf Gradient and Temperature Response in Barley

**DOI:** 10.1101/2025.06.24.661333

**Authors:** Edward Cedrick Fernandez, Gaoya Tu, Wei Dai, Shengming Yang, Zhaohui Liu, Marcin Grzybowski, Zhikai Liang

**Author notes:** **Corresponding author:** Zhikai Liang.

## Abstract

The linear shape of cereal leaves creates distinct longitudinal zones that coordinate tissue maturation and resource allocation. Under abiotic stress such as heat, these longitudinal zones may differentially activate protective pathways, revealing hidden heterogeneity in stress response that remains poorly understood. Barley (*Hordeum vulgare*), a cold-adapted crop particularly sensitive to elevated temperatures, can serve as an ideal model for studying region-specific heat responses in leaves. Using chlorophyll fluorescence imaging, we found that non-photochemical quenching (NPQ) kinetics captured additional physiological changes beyond those detected by SPAD, highlighting the added value of chlorophyll fluorescence-based assessments. NPQ kinetics traits displayed consistent spatial gradients from tip to base, with heat stress reducing NPQ induction across leaf gradients. Genome-wide association analysis of traits derived from chlorophyll fluorescence imaging across leaf gradients identified significant SNPs within multiple candidate genes, including *HORVU.MOREX.r3.3HG0262630* that was consistently detected in over 90% resampling iterations under heat stress, suggesting its key role during heat responses. Transcriptomic profiling along the leaf axis revealed both conserved and region-specific heat responses between leaf regions. Conserved activations highlighted conserved heat response pathways, including the reactivation of the *Arabidopsis* thermomemory module *FtsH6-HSP21*. In contrast, the region-by-temperature interaction analysis identified 40 genes with spatial responses indicative of resource reallocation from growth to defense, including those involved in growth and transport. The integration between spatially resolved phenotyping and transcriptional profiling underscores region-specific variation in response to heat along the leaf axis, guiding targeted strategies to enhance heat resilience in barley and other cereal crops.

## Introduction

Barley (*Hordeum vulgare*) is a temperate cereal adapted to cool climates and is vulnerable to elevated temperatures during critical growth stages (Jacott and Boden, 2020). As a major abiotic factor, high temperature can lead to collapse of intracellular organization, disruption of cell membrane integrity and suppression of photosynthetic function, ultimately impacting agricultural performance like grain filling, harvest index and tillering (Barnabás *et al*., 2008; Allakhverdiev *et al*., 2008). Of these negative effects, the destabilization of photosystem II (PSII) – a core component of the photosynthetic machinery – is especially detrimental. Under environmental stresses like high temperature, photosynthetic efficiency is often compromised due to chlorophyll degradation, destabilization of PSII reaction centers, and impaired excitation energy transfer within the thylakoid membrane, disrupting the capacity of plants for effective carbon assimilation (Allakhverdiev *et al*., 2008). Improving photosynthetic efficiency has demonstrated strong potential to enhance fitness across a range of plant species (Feng *et al*., 2014; De Souza *et al*., 2022; Qu *et al*., 2021; Li *et al*., 2020; Athanasiou *et al*., 2010), emphasizing the need to elucidate the genetic mechanisms that govern photosynthetic resilience under high temperature conditions.

Chlorophyll fluorescence imaging provides a robust quantitative approach for assessing PSII efficiency and evaluating stress tolerance in plants (Guidi *et al*., 2019). When applied at the population scale, this approach enabled the identification of key photosynthesis-related genes – such as PSII subunit S (*PSBS*) (Sahay *et al*., 2023; Rungrat *et al*., 2016; Wang *et al*., 2017; Sahay *et al*., 2024) – that regulate photoprotection and energy dissipation. Despite this potential, the environmental plasticity of genetic loci governing photosynthetic responses – particularly in parallel, well-controlled abiotic stress environments – remains uncharacterized. This gap limits our understanding of how genotypes dynamically adjust photosynthetic performance across contrasting environmental conditions. Among chlorophyll fluorescence traits, Non-Photochemical Quenching (NPQ) functions as a critical photoprotective mechanism under stress environments by dissipating excess excitation energy as heat, thereby safeguarding PSII from photodamage. Due to its central role in regulating energy balance within the chloroplast, variation of NPQ across genotypes holds promise as an effective indicator for assessing heat tolerance in a natural population.

In fact, photosynthesis may not be uniformly regulated across different regions of a leaf. Developmental gradients, such as those from the base to the tip, give rise to spatial heterogeneity in cellular structure and chloroplast function (Schurr *et al*., 2006). In parallel, region-specific microenvironments can influence local light interception, hormone signaling and molecular pathways, ultimately leading to differential photosynthetic performance along the leaf axis (Fukshansky and Remisowsky, 1992). At the cellular level, chloroplasts can actively reposition in response to light conditions to optimize energy capture and minimize photodamage, contributing to spatial heterogeneity in photosynthetic efficiency even within the same tissue (Kasahara *et al*., 2002). Although gene expression varies along the developmental gradient in both C4 and C3 plant leaves (Li *et al*., 2010; Li *et al*., 2015; Nelson, 2011), the genetic mechanisms underlying region-specific responses to abiotic stress within a single leaf remain largely unresolved. High-resolution fluorescence imaging has shown that intra-cellular variability in stress responses – such as differences in PSII efficiency and NPQ among chloroplast clusters – can be particularly pronounced under environmental stress (Sekulska-Nalewajko *et al*., 2019; Pérez-Bueno *et al*., 2019; Herritt *et al*., 2020). Despite these insights, the underlying regulatory mechanisms driving such spatial heterogeneity are still largely uncharacterized. Some photoreceptor and chloroplast movement genes, PHOT1 (Liscum and Briggs, 1995) and CHUP1 (Oikawa *et al*., 2008), have been implicated in photochemical response dynamics, but it is still unclear how genetic and transcriptional programs contribute to the spatial tuning of photosynthesis within a tissue. Uncovering region-specific genes that regulate spatial variation in photosynthetic plasticity under stress could provide new insights into the mechanisms of plant resilience. Such knowledge has important implications for crops like barley and other cereals, where localized stress responses along the leaf may significantly impact overall physiological performance.

The barley leaf exhibits a clear developmental gradient from the proliferative base to the mature tip, creating spatial zones with distinct cellular identities and physiological capacities. This intrinsic gradient offers a framework to investigate how environmental signals interact with developmental programs to shape localized stress responses – coordination that is critical for sustaining photosynthetic efficiency and growth under heat stress. While genotypic variation in heat tolerance is well-documented, how different regions within a single leaf differentially perceive and respond to thermal stress remains poorly understood. Exploring this spatial interplay between development and stress signaling may reveal how plants fine tune resilience at the regional level.

In this study, we leveraged a subset of 160 selected diverse spring barley accessions from the International Barley Core Collection (BCC) (Muñoz-Amatriaín *et al*., 2014) and exposed them to paired control and heat stress conditions. By integrating population-scale regional chlorophyll fluorescence imaging with transcriptomic profiling of a reference genotype, we revealed coordinated phenotypic and molecular responses across defined regions along the developmental gradient at the individual leaf level. This spatially resolved approach enabled the identification of region-specific genetic factors underlying heat stress responses within the leaf, providing insights for breeding heat-tolerant barley and advancing the understanding of thermal resilience in crops.

## Results

### Spatial Variations in Leaf Chlorophyll Content Under Contrasting Temperature Regimes

To assess the impact of elevated temperature on barley, we evaluated 159 lines from a previously curated panel of ∼200 genotypes (Karki *et al*., 2024), originally selected from the Barley Core Collection (BCC) (Muñoz-Amatriaín *et al*., 2014) using molecular markers to capture the broad genetic diversity, with the genotype Morex included as the check line. At 10 days after planting (DAP), plants were moved from the greenhouse to growth chambers and subjected to either a 3-day heat treatment (33°C/26°C day/night) or maintained 3-day under a control condition (23°C/16°C day/night). Chlorophyll content was estimated for each plant at the end of the treatment at DAP 14 using a Soil Plant Analysis Development (SPAD) 502 chlorophyll meter. This experiment was separately repeated five times (see Methods), resulting in approximately 650 individual plants available for analysis in each condition.

Because barley leaves have a linear shape and develop progressively from base to tip, we anticipated that different developmental regions along the leaf might exhibit variable responses to temperature. To capture this spatial variation, we measured three distinct regions – base, middle, and tip – separately along each leaf (Figure 1a). Among the accessions evaluated, the highest average SPAD values were recorded in PI 329032 (48.82) under control conditions and PI 611555 (45.39) under heat conditions. In contrast, the lowest values were observed in PI 190790 (28.42) and CIho 14291 (26.76) under control and heat stress conditions, respectively. However, SPAD values varied substantially across leaf regions based on measurements collected from different regions (Supplementary Table 1). To quantify this spatial variation, we calculated the coefficient of variation of SPAD values across three leaf regions, enabling standardized comparisons of variability along the leaf gradient, even in the presence of large differences in mean values. The broad-sense heritability of the coefficient of variation was then estimated at ∼0.604 under both conditions, indicating that spatial variation in SPAD values was partially controlled by genetic factors. In addition, coefficients of variations were highly correlated between two tested temperature regimes, suggesting that the pattern of spatial variability in SPAD values was also largely conserved across environments (Pearson r = 0.497, p-value = 2.71e^−11^) (Supplementary Figure 1).

**Figure 1.**
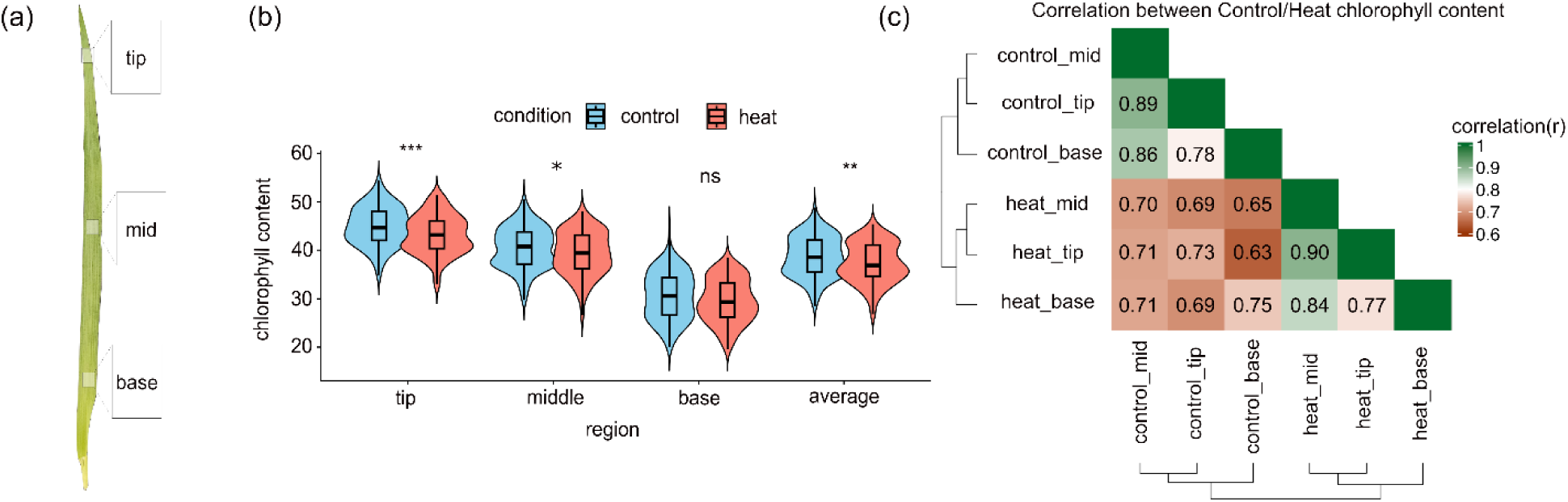
Spatial gradient of chlorophyll responses to heat stress along barley leaves. (a) Each leaf per barley plant was sampled from three leaf regions: tip, middle and base; (b) Chlorophyll content estimated by SPAD across leaf regions under control and heat stress conditions. Each data point represents the mean chlorophyll content across all plants within that gradient for a given accession. Asterisks denote significance levels (*** p < 0.001, ** p < 0.01, * p < 0.05; ns = not significant); (c) Pairwise Spearman correlations were calculated among the three leaf regions to assess the consistenc of chlorophyll content patterns under control and heat conditions. BLUEs of chlorophyll content per accession and leaf region were used for these analyses. Hierarchical clustering was applied to group leaf regions and conditions.

Under elevated temperature, chlorophyll content was significantly reduced across the population, with the most significant decline observed at the leaf tip (paired Wilcoxon test; p-value = 2.28e^−4^) (Figure 1b). This negative effect progressively diminished from the tip toward the base of the leaf (Figure 1b). After accounting for covariate effects, we calculated the Best Linear Unbiased Estimates (BLUEs) of SPAD values for each accession and examined the correlation between BLUEs under control and heat conditions. Under both conditions, SPAD BLUEs exhibited strong correlations (Spearman’s r > 0.845, p-value < 2.23e^−308^) between the middle region and either the tip or base, whereas correlations between the tip and base were obviously lower. However, the correlation coefficient decreased to approximately 0.700 between control and heat conditions (Figure 1c).

### Assessing Photosynthetic Efficiency Using Chlorophyll Fluorescence Imaging

While SPAD offers a rapid estimate of chlorophyll content in accessing plant health, it does not capture the dynamic photochemical responses of plants to stress. To evaluate the impact of elevated temperature on barley, we employed the PlantExplorer PRO+ chlorophyll fluorescence imaging system (PhenoVation, Netherlands) to extract NPQ, chlorophyll index (ChlIdx), anthocyanin index (AriIdx) and normalized difference vegetation index (NDVI) from fully expanded leaf images in the same panel used for SPAD measurements. Building on a previously established approach (Sahay *et al*., 2024), exponential and hyperbolic models were fitted to NPQ kinetics curves, covering both induction and relaxation phases. Thi enabled us to extract 14 quantitative NPQ kinetics traits that captured the dynamics, steady-state levels, and amplitude of photosynthetic energy dissipation responses. Together, these measurements from chlorophyll fluorescence images yielded 17 spatially resolved traits describing photoprotective responses, pigment composition, and vegetation vigor (Figure 2a; Supplementary Table 2).

**Figure 2.**
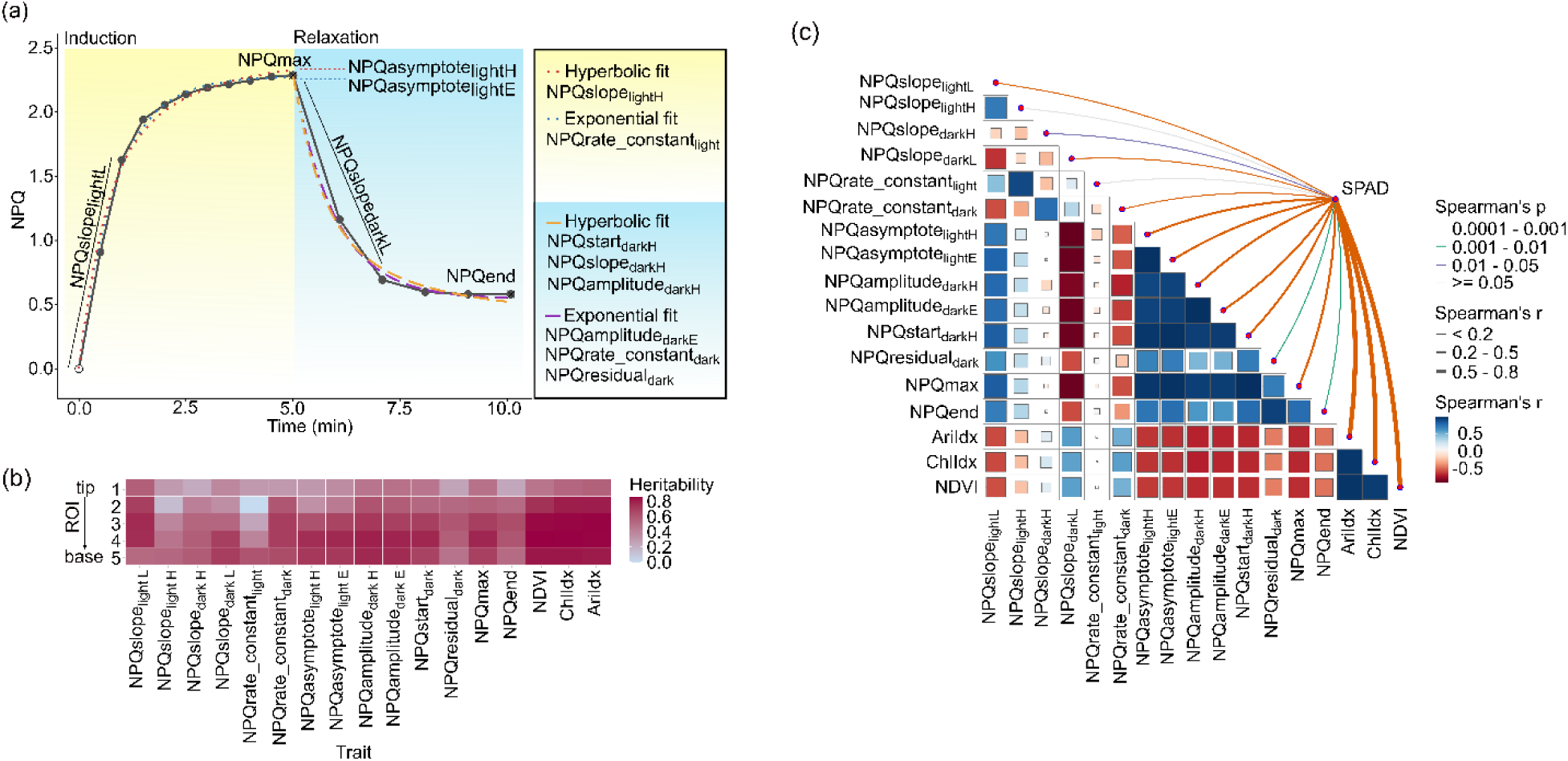
Non-photochemical quenching (NPQ) kinetics traits for evaluating photosynthetic efficiencies. (a) An example of the NPQ curve was generated using a sample from ‘Morex’, specifically at ROI 4 under the heat stress condition. Individual NPQ kinetics traits were derived by fitting hyperboli and exponential models applied to the NPQ curve using Equations 1 to 4 that were previously defined (Sahay *et al*., 2024). Data collection began after 30 seconds of illumination; (b) Broad-sense heritabilit estimates for each NPQ kinetics trait across five ROIs (Regions of Interest) from tip to base under heat stress condition (ChlIdx: Chlorophyll index; AriIdx: Anthocyanin Index; NDVI: Normalized Difference Vegetation Index); (c) Spearman’s rank correlation matrix illustrates the relationships between NPQ kinetics traits and SPAD values under heat stress conditions. Both traits from chlorophyll fluorescence images and SPAD measurements were obtained from the same leaf, specifically targeting the middle region of each plant. The correlation matrix was calculated using BLUE values derived from the middle leaf regions of 159 accessions.

To enable spatially resolved analysis of phenotypic responses to temperature, all 17 traits from five evenly spaced regions of interests (ROIs) spanning from tip to base of an individual leaf were extracted (Supplementary Figure 2). Assessment of individual traits in this panel revealed that broad-sense heritability (H²) exhibited a gradient decline from the leaf base to the tip for most traits (Figure 2b; Supplementary Figure 3). With the exception of traits measured in the leaf tip region, ChlIdx, AriIdx, and NDVI consistently showed higher H² compared to NPQ kinetics traits under both control and heat conditions (Figure 2b; Supplementary Figure 3; Supplementary Table 3). Among NPQ kinetics traits, those associated with response rate (*e.g.*, NPQrate_constant_light_) displayed lower heritability compared to traits representing steady-state responses (*e.g.*, NPQ_max_) (Figure 2b). For example, across all five ROIs, the average heritability of NPQrate_constant_light_ was 0.309 while that of NPQ_max_ average heritability was 0.638.

To compare the correlation between SPAD values and all 17 traits derived from chlorophyll fluorescence images, we collected both SPAD measurements and fluorescence images from the same leaf of each plant grown under heat conditions. A total of 650 individual leaves, representing 159 accessions across five experimental replicates, were retained for analysis, with multiple Morex plant included to control for batch effects. After accounting for experimental covariate effects, we calculated the BLUEs for SPAD and each of 17 traits from chlorophyll fluorescence images at the middle region of the leaf for each accession (Supplementary Table 4). AriIdx, ChlIdx, and NDVI were highly correlated with one another (r > 0.953; p-value < 3.72e^−83^), demonstrating strong consistency among these multispectral indices. Interestingly, SPAD values were also highly correlated with all three indices – AriIdx, ChlIdx, and NDVI (r > 0.833; p-value < 3.79e^−42^), suggesting overall alignment while potentially capturing distinct yet complementary aspects of leaf optical properties (Figure 2c). In contrast, NPQ kinetics traits, derived from time-resolved chlorophyll fluorescence imaging, represent a distinct class of dynamic measurements that quantified the regulation of excess energy dissipation through non-photochemical quenching in PSII. NPQasymptote_lightH_ and NPQasymptote_lightE_, derived from curve fitting during the induction phase of NPQ kinetics, were both strongly correlated (r > 0.971, p-value < 8.20e^−100^) with NPQ_max_ – the peak non-photochemical quenching value attained during induction, reflecting their shared dependence on the dynamic accumulation of energy dissipation capacity. The lack of strong correlation between NPQ kinetics and multispectral indices such as ChlIdx and AriIdx (absolute r value ranged from 0.005 to 0.711) also highlighted their complementary roles in assessing photosynthetic performance (Figure 2c).

### Dynamic Variations of NPQ Kinetics between Contrasting Temperature Conditions

To evaluate spatial variation in temperature responses across individual barley leaves, chlorophyll fluorescence images were collected from 633 and 650 leaves under control and heat conditions, respectively, encompassing all five experimental replicates. This resulted in the analysis of over 6,000 individual leaf regions. Instead of focusing on a specific position of each individual leaf, we captured the entire leaf to assess spatial variation of photosynthetic efficiency along the leaf gradient (Supplementary Figure 2). For NPQ kinetics analysis, five ROIs from base to tip were evenly selected from the chlorophyll fluorescence image of each leaf. A gradient change of NPQ kinetics traits was observed from tip (ROI 1) to tip (ROI 5). Specifically, NPQ_max_ progressively increased from the leaf tip to the base, with the basal region showing the highest NPQ_max_ and the tip region having the lowest NPQ_max_. This pattern highlighted a spatial gradient in energy dissipation capacity along the leaf axis in barley in either temperature regime (Figure 3a). The slope of NPQ induction increased from tip to base, suggesting faster activations of photoprotective mechanisms. The reduction of NPQ_end_ from control to heat was smallest at the leaf tip (ROI 1; Δ = 0.187) and largest at the base (ROI 5; Δ = 0.365). This difference suggested that basal leaf regions, despite having lower chlorophyll content, experienced a greater disruption in energy dissipation under heat stress.

**Figure 3.**
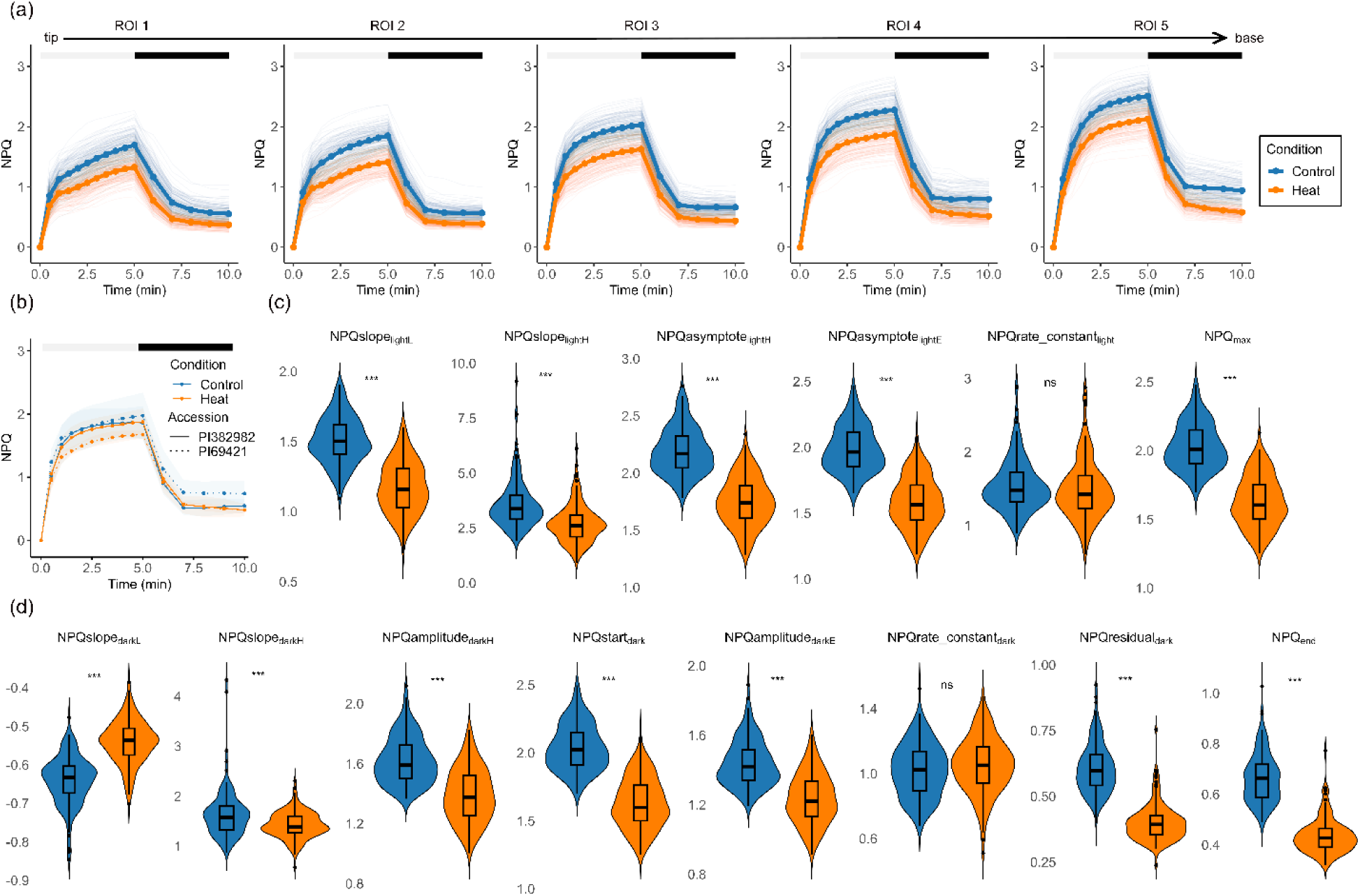
NPQ responses under control and heat stress conditions. (a) NPQ dynamics during the induction phase (white bar) and relaxation phase (black bar) across five defined regions of interest (ROI 1-5) from leaf tip to base under control and heat stress conditions (n = 159). Each curve represents the mean NPQ response of all plants for each genotype and treatment condition; (b) Representative NPQ response curves from middle regions of leaves (ROI 3) under control and heat stress conditions for two contrasting genotypes. The number of biological replicates per genotype is 5; (c) Induction phase trait and (d) relaxation phase traits extracted from the middle region of leaf (ROI 3) for all genotypes under control and heat stress conditions. Statistical significance is indicated as: ***p < 0.001, **p < 0.01, *p < 0.05, and ns = not significant.

Of the 159 barley accessions studied, NPQ responses to heat stress varied widely. Focusing on the central leaf region (ROI 3), accession PI 382982 exhibited minimal changes in NPQ kinetics between control and heat conditions. In contrast, PI 69421 showed a pronounced reduction in NPQ under heat stress, indicative of a distinct photoprotective strategy (Figure 3b). Interestingly, PI 69421 originated from the North of China and PI 382982 from Ethiopia (from USDA GRIN), highlighting the potential influence of geographical and latitude origins on temperature response patterns. Of 14 NPQ kinetics traits, 6 traits were calculated under the light phase and 8 traits were calculated under the dark phase. Light-phase NPQ kinetics traits, including NPQ_max_, significantly decreased (paired Wilcoxon test; p-value < 0.001) under heat, suggesting a reduced photoprotective response during illumination (Figure 3c). Interestingly, F_v_/F_m_ values were elevated under heat stress compared to control conditions. While the underlying mechanism remained unclear, this observation might reflect modulation of PSII efficiency during early or moderate stress exposure (Supplementary Table 2). Most dark-phase NPQ kinetics traits declined under heat conditions compared to control, reflecting an overall decrease in quenching capacity. However, NPQslope_darkL_ increased under heat, suggesting a slower initial rate of NPQ relaxation in heat than control after the light was turned off (Figure 3d).

### Dissecting genetic signals in related to the response to heat stress

To identify genetic loci associated with all 17 traits derived from chlorophyll fluorescence images acros temperature regimes, we conducted a genome-wide association study (GWAS) using the resample model inclusion probability (RMIP) approach. A total of 14 NPQ kinetic traits, along with three additional multispectral indices – AriIdx, ChlIdx and NDVI – were analyzed across five longitudinal leaf regions spanning from base to tip under both control and heat stress conditions. In total, 17 traits were analyzed in GWAS using 6,574 previously characterized high-quality Single Nucleotide Polymorphism (SNPs) (Morales *et al*., 2022). We identified 32 significant (RMIP threshold > 0.500) SNP-trait associations using this approach. All associated SNPs mapped directly to 13 genes (0 bp distance to the corresponding SNPs), with 3 identified under control conditions and 10 under heat stress (Figure 4; Supplementar Figure 4; Supplementary Table 5). The strongest RMIP signal driven by SNP *11_10328* was repeatedl detected in 185 out of 200 resampling iterations (RMIP = 0.925) for NPQrate_constant_dark_ in ROI 4 under the heat condition, while this SNP consistently exhibited high RMIP (RMIP > 0.500) values across all regions except ROI 1. By running the GWAS using all 159 samples, we were still able to identify the significant association (p-value = 3.86e^−8^) between NPQrate_constant_dark_ and SNP *11_10328* (Supplementary Figure 5a). Inbred lines with the reference allele and inbred lines with the alternative allele showed internally consistent NPQ rate constant values – but these balanced values significantly differed between the two allele groups across the leaf gradient except at ROI 1 (Supplementary Figure 5b). This SNP *11_10328* was located at 266,345,990 bp of chromosome 3H within the gene *HORVU.MOREX.r3.3HG0262630*, specifically on the third exon that led to a missense mutation. Orthologs of *HORVU.MOREX.r3.3HG0262630* in *Arabidopsis* encode for secretion-associated RAS (SAR) superfamily proteins, part of the small ADP-ribosylation factors (ARF)-like GTPase family involved in vesicle formation and trafficking between the endoplasmic reticulum and Golgi apparatus. These proteins are essential for cellular homeostasis and rapid stress responses (Brandizzi, 2018). Several studies revealed that SAR/ARF was increasingly induced under both abiotic and biotic stress conditions, including wheat (Li *et al*., 2021), rice (Muthamilarasan *et al*., 2016; Joshi *et al*., 2014) and spinach (Zhao *et al*., 2018). The leaf gradient-based GWAS offered the potential to enhance signal detection by reducing noise and resolving region-specific association signals. For example, SNP *SCRI_RS_223734* (RMIP = 0.560) located within *HORVU.MOREX.r3.4HG0403260* was strongly associated with NPQrate_constant_dark_ only in ROI 4 under heat conditions, but not in other regions. The *Arabidopsis* ortholog *AT5G61410* encoding D-ribulose-5-phosphate-3-epimerase (RPE) plays a role in both the Calvin–Benson cycle and the oxidative pentose phosphate pathway. *Arabidopsis rpe* mutants with reduced RPE abundance have shown elevated steady-state NPQ levels, indicating impaired photosynthetic electron transport (Li *et al*., 2022). In addition, Cold stress-induced proteomic changes in *Arabidopsis* chloroplasts revealed reduced RPE abundance, pointing to a potential role for RPE in abiotic stress adaptation (Goulas *et al*., 2006). Under heat stress, SNP *12_31059* emerged as a potential pleiotropic locus within *HORVU.MOREX.r3.5HG0511430*, exhibiting significant associations (RMIP > 0.535) with nine NPQ kinetics traits. *HORVU.MOREX.r3.5HG0511430* encodes a protein with WD40 repeat domains implicated in environmental adaptation in other species (Eryong and Bo, 2022; Kong *et al*., 2015; Liu *et al*., 2017; Lee *et al*., 2010; Xu *et al*., 2019). The RMIP value of SNP *12_31059* also differed by more than 0.500 between heat and control conditions, indicating a potential genotype-by-environment interaction. In total, five SNP-trait associations with RMIP differences ≥ 0.500 were identified across environments, all linked to two SNPs – SNP *12_31059* and *SCRI_RS_126369* (Supplementary Table 5). The other SNP, *SCRI_RS_126369*, was associated with NPQslope_lightH_ and NPQslope_lightL_ in ROI 3 and was located within *HORVU.MOREX.r3.3HG0323480* that encodes a RRM (RNA recognition motif) domain-containing protein. In *Arabidopsis*, loss-of-function mutations in the homologous gene *AT1G73530* (encoding ORGANELLE RNA RECOGNITION MOTIF PROTEIN 6, ORRM6) caused a 41-51% reduction in NPQ, indicating impaired PSII efficiency and photoprotective capacity, and suggesting a potential functional role of *HORVU.MOREX.r3.3HG0323480* in barley (Hackett *et al*., 2017).

**Figure 4.**
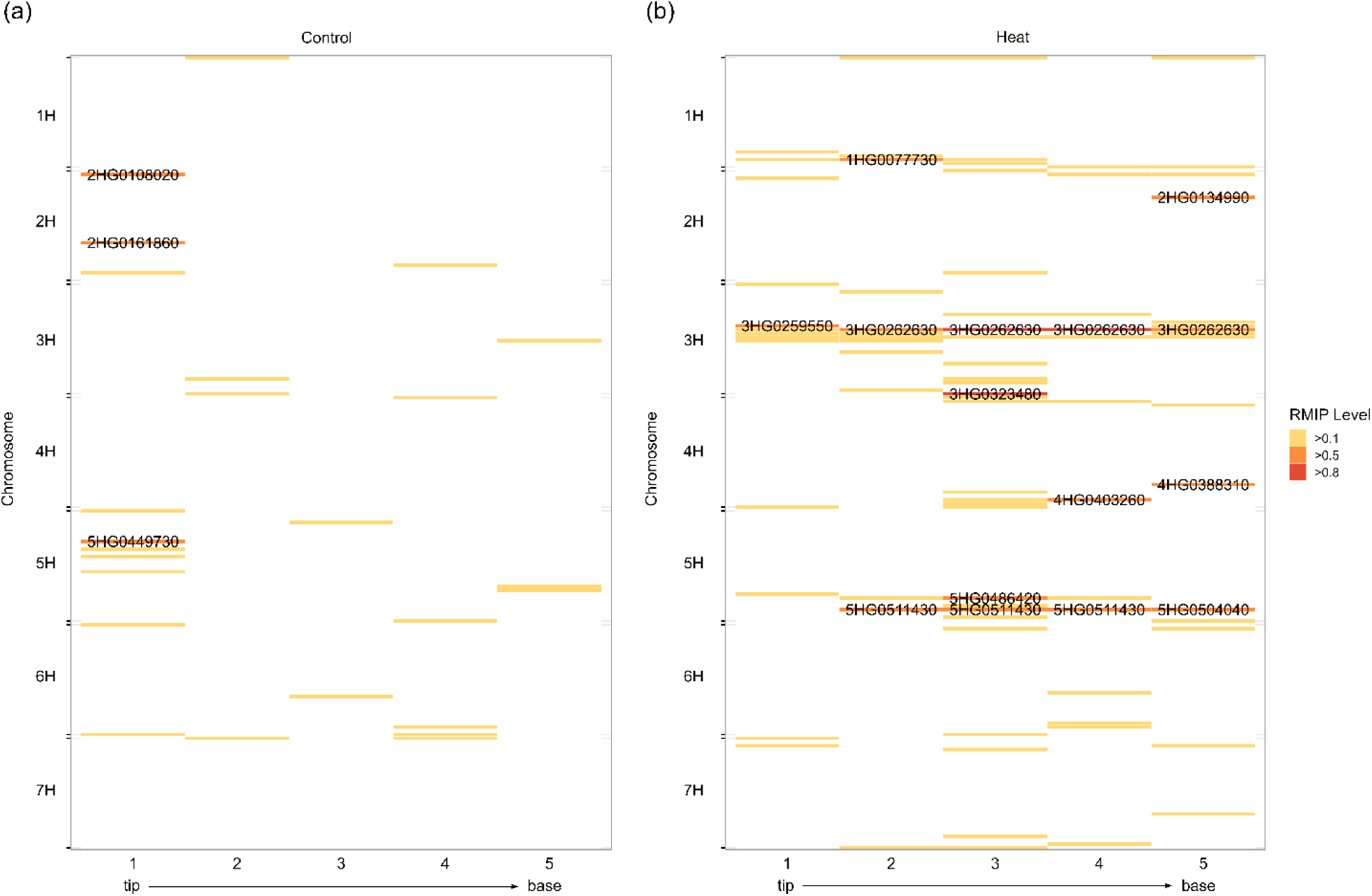
RMIP-GWAS summary for chlorophyll fluorescence traits under control and heat conditions. RMIP-based GWAS was conducted for each of 17 traits across five equally spaced leaf regions, ranging from tip to base. Each chromosome was partitioned into 30 equal-sized bins. For each bin, the SNP with the highest RMIP value relative to any studied traits was selected to represent that region. Bins with RMIP values ≤ 0.100 are shown in white. Genes located near high-RMIP SNPs are annotated within the colored bars. Each bin with the RMIP > 0.500 was annotated with the respective gene name, and the ‘HORVU.MOREX’ prefix was removed from all gene IDs for easy visualizations. (a) Control condition; (b) Heat condition.

### Transcriptomic response to heat stress across leaf gradients

While GWAS provided a population-level understanding of genetic loci associated with phenotypi variation under heat stress, transcriptome profiling will offer a mechanistic perspective on how plant respond to stress at the molecular level. Integrating both approaches may reflect the distinct yet complementary roles of inherited genetic variation and dynamic gene regulation in shaping heat responses across leaf gradients. Using the barley reference genotype Morex, we collected base, middle, and tip regions from fully expanded leaves of plants exposed to a 4-hour heat stress or control conditions to generate RNA-seq data. After removing genes with no counts across all samples, we obtained 22,482 genes that were present in all samples and 14,218 of them were expressed with mean CPM (Counts Per Million) > 1. Samples from the base and tip regions of the leaf were clearly separated, whereas those from the middle region showed partial overlap with the base samples (Supplementary Figure 6), reflecting their transitional role between the two regions. In total, we identified 675, 866, and 1,419 differentiall expressed genes (DEGs) between control and heat conditions in the base, middle, and tip regions, respectively, with 372 commonly shared across all three leaf regions (Figure 5a). The GWAS-identified SNP (RMIP = 0.595 under heat stress), located within *HORVU.MOREX.r3.5HG0511430* and potentially pleiotropic, was not classified as a DEG but nonetheless showed moderate induction by heat stress across all three leaf regions, with consistently high baseline expression under both control and heat conditions (mean CPM under control = 15.17; mean CPM under heat = 20.93; log_₂_ fold change = 0.618). By focusing on genes with a log2 fold change (Heat/Control) > 5 in both the leaf base and tip – two regions representing distinct developmental zones – we identified 21 shared DEGs, 71% of which were related to heat shock proteins (HSPs). This observation recapitalized that the high-regulation pattern of HSPs was not only conserved across genotypes – as previously observed in a maize population (Liang *et al*., 2022) but also conserved across anatomically distinct regions along the leaf axis. The most upregulated gene, *HSP21* (*HORVU.MOREX.r3.4HG0392240*), was identified to protect PSII during exposure to elevated temperatures (Neta-Sharir *et al*., 2005). Interestingly, we also observed significant co-upregulation across all three leaf regions of the FtsH6-HSP21 regulatory module that was shown to mediate thermomemory in *Arabidopsis* (Sedaghatmehr *et al.,* 2016) (log2 fold change > 5, adjusted p-value < 0.05; Figure 5b). However, none of the region-specific DEGs were associated with putative HSP functions. Instead, some downregulated region-specific genes might play roles in coordinating the interplay between cellular development and heat stress responses. For instance, *HORVU.MOREX.r3.2HG0113890*, the barley ortholog of *Arabidopsis* GH3.9 (*AT2G47750*), was specifically downregulated in the leaf base under the heat condition (Supplementary Table 6). Given GH3.9 genes were found to typically promote growth under non-stress conditions (Khan and Stone, 2007), its suppression in barley might indicate a shift in resource allocation from growth to defense.

**Figure 5.**
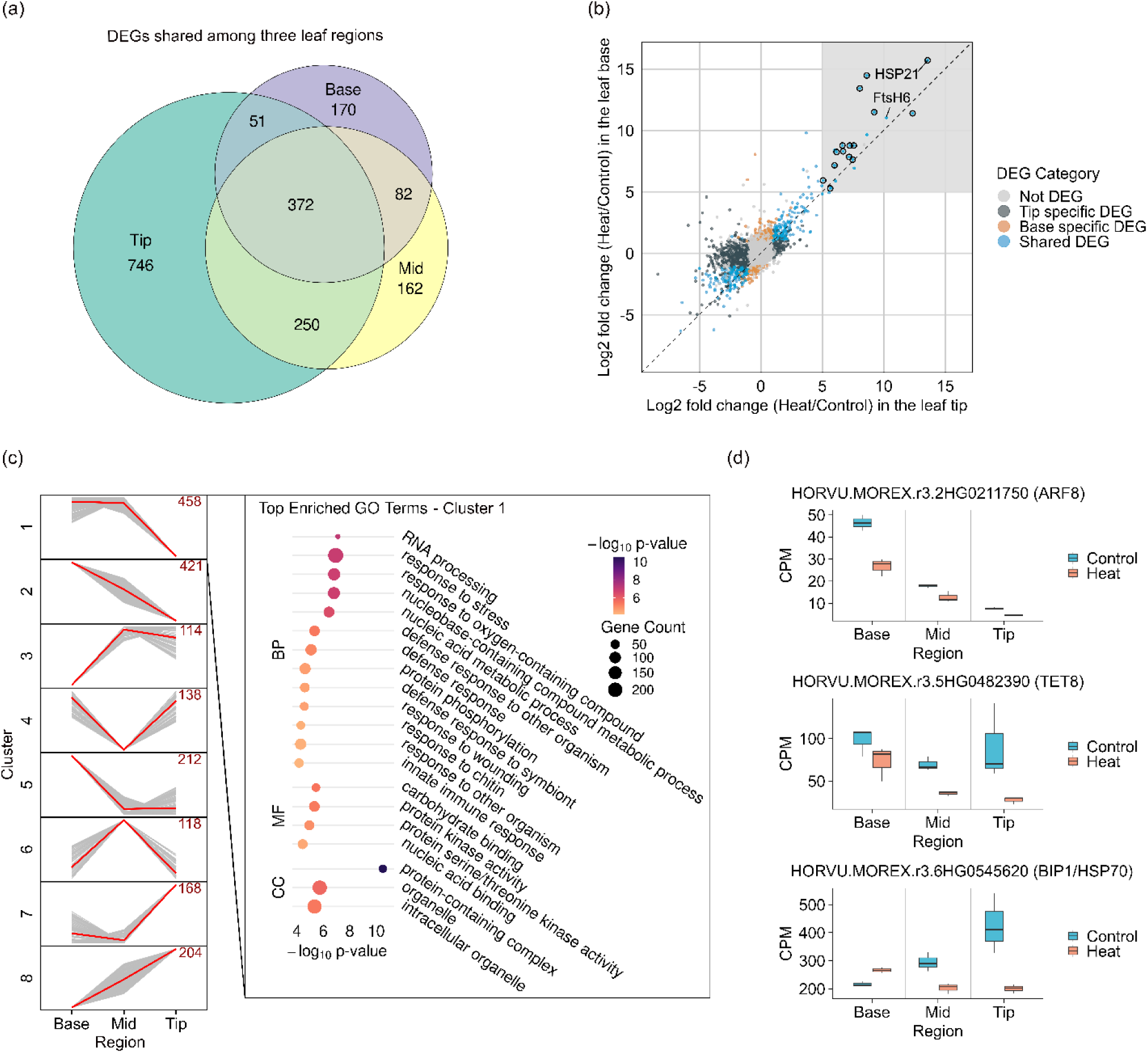
Spatially resolved transcriptomic responses to heat stress along the leaf gradient in barley. (a) Overlap of DEGs among the base, middle, and tip regions collected from the same leaves of the barley genotype Morex under heat stress conditions; (b) Comparison of gene responses between tip and base regions under heat stress, based on log_₂_ fold change values. A customized threshold as log_₂_ fold change > 5 is used to define genes as highly upregulated in both base and tip regions. Genes with putative HSP functions are highlighted in black circles. The previously characterized FtsH6-HSP21 regulatory module is annotated; (c) Classification of DEGs into eight expression clusters using k-mean clustering of normalized log_₂_ fold changes. The number of genes per cluster is indicated. Functional GO enrichment of genes in the cluster is highlighted for Gene Ontology biological process (BP), molecular function (MF), and cellular component (CC) terms. P-values were corrected using the Bonferroni method; (d) Three examples of genes identified as differentially expressed or showing significant region-by-temperature interactions were selected to show. Gene identities are based on *Arabidopsis thaliana* orthologs annotated by *Ensembl Plants*.

To assess transcriptional responses to heat stress across the leaf gradient in barley, a union set of DEGs from the base, middle, and tip regions was compiled to enable comprehensive clustering of heat response patterns. These genes were then grouped into eight distinct clusters using k-means clustering across three leaf regions. Of the eight clusters, only clusters 1, 2 and 5, which exhibited relatively decreasing response patterns from base to tip, showed enriched GO (Gene Ontology) terms, many of which were related to environmental stress responses (*e.g*., GO:0050896) (Figure 5c; Supplementary Table 7). However, all other clusters had only less than two enriched GO terms. These clustering patterns reflected broader trends in gene responses to heat stress across the leaf gradient. For example, we identified an auxin response factor (ARF), *HORVU.MOREX.r3.2HG0211750*, as a representative gene associated with growth regulation. Its expression consistently declined from the leaf base to the tip under both control and heat conditions (Figure 5d). The further reduction in expression under heat stress suggests a shift away from developmental prioritization, potentially reflecting a trade-off between growth and stress adaptation. In contrast, region-specific signals prioritized overlaps between GWAS and RNA-seq data. For example, SNP *12_20297*, located within *HORVU.MOREX.r3.5HG0482390* that encodes a tetraspanin family protein, was associated with both ChlIdx (RMIP = 0.435) and AriIdx (RMIP = 0.420) in the GWAS for leaf tip traits under heat stress, and this gene was significantly downregulated at the leaf tip in RNA-seq data (Figure 5d). However, the magnitude of this downregulation was smaller in the base and middle regions of the leaf. By leveraging a linear model to identify genes with significant interaction effects between temperature and base-tip comparisons, we identified 40 genes that were differentially regulated in response to heat stress between leaf regions (adjusted p-value < 0.001 (Zhang *et al*., 2017)). This analysis of leaf gradient responses uncovered regulatory patterns that might have been overlooked. Although most HSP displayed conserved upregulation patterns, *HORVU.MOREX.r3.6HG0545620*, encoding a putative HSP was identified to be slightly upregulated (log2 fold change = 0.306) in the base but downregulated in other regions (log2 fold change in middle as -0.537, tip as -1.081) (Figure 5d).

## Discussion

As a cool-season crop, barley has low tolerance to elevated temperatures. Specifically, daytime temperatures in the range of 30-36°C were commonly used to simulate heat stress in barley (Sakata *et al*., 2010; Ingvordsen *et al*., 2018; Mahalingam and Bregitzer, 2019). In this study, heat stress was applied as a +10°C increase above the control temperature, using two distinct durations: a 3-day treatment (33°C/16°C day/night) for phenotyping and a 4-hour exposure (33°C) for RNA-seq analysis.

The 3-day treatment was necessary to elicit measurable changes in physiological traits such as NPQ dynamics and chlorophyll indices, which require time to accumulate and reflect integrated stress outcomes. Prior studies have identified that abiotic stress including heat stress can decrease NPQ values compared to control environments (Zhou *et al*., 2017; Zhou *et al*., 2018; Lauterberg *et al*., 2024; Qu *et al*., 2025). A previous study found that NPQ decreased more under heat stress than under drought or combined drought and heat stress in tomato (Zhou *et al*., 2017; Zhou *et al*., 2018; Lauterberg *et al*., 2024; Qu *et al*., 2025), suggesting that elevated temperature may exert a more direct or acute disruption to PSII function and photoprotective capacity. Consistent with previous findings, our results also showed that the overall mean NPQ decreased under elevated temperature conditions compared to the control, and most of 14 derived NPQ kinetics traits show significant differences between control and heat conditions (Figure 3a). While NPQ reflects the activation of photoprotective energy dissipation, F_v_/F_m_ may serve as a direct indicator of PSII photochemical efficiency. However, the utility of F_v_/F_m_ in capturing PSII alterations under heat stress may be limited, as it often remains relatively stable or less sensitive to moderate increases in temperature (Crafts-Brandner and Salvucci, 2002). A previous study revealed prolonged heat stress typically reduces F_v_/F_m_ in plants (Murchie and Lawson, 2013), but others also reported increases in F_v_/F_m_ following short-term heat exposure (Zhou *et al*., 2018; Crafts-Brandner and Salvucci, 2002; Wen *et al*., 2005), potentially reflecting adaptive or protective responses. However, population-scale evaluations comparing F_v_/F_m_ responses across contrasting temperature conditions in crop species remain limited, highlighting the need for broader investigations into genotype-by-environment interactions affecting PSII photochemical efficiency. Interestingly, our study identified a ∼3% increase of mean F_v_/F_m_ values across all leaf regions under heat (Supplementary Table 2) compared to control, suggesting that photoprotective mechanisms might not have been fully engaged despite underlying heat stress. The observed reduction in NPQ alongside a slight increase in F_v_/F_m_ might indicate a transient decline in PSII efficiency under heat stress, consistent with the hypothesis that elevated temperatures might impair PSII function without causing permanent structural damage to the photosynthetic apparatus (Mathur *et al*., 2014).

The effect of our simulated heat stress was validated through RNA-seq analysis of the barley genotype Morex exposed to a 4-hour 33°C in the daytime. This 4-hour treatment was selected to capture early, transient transcriptional changes that reflect immediate stress perception and signal initiation (Kollist *et al*., 2019), before widespread reprogramming or damage responses obscure the primary regulatory signals. Specifically, we identified 31 putative HSPs that were commonly upregulated in all regions, consistent with known activation patterns of HSPs under heat conditions (Vierling, 1991). To better disentangle these signals between widespread transcriptional reprogramming under acute heat stress and phenotypic adaptation under moderate heat exposures, we integrated GWAS and RNA-seq datasets to pinpoint genetic loci linked to transcriptomic responses. Using this approach, 10 DEGs were identified that either showed significant expression differences between conditions or significant region x temperature interaction effects and overlapped with at least one GWAS hit detected ≥ 10% of resampling iterations. These findings highlighted both transcriptional responsiveness and genetic co-localization with phenotypic variation, suggesting potential causal roles in heat stress adaptation. While the current overlap may be constrained by marker resolution, future improvements in genotyping density may enhance the detection of regulatory variants linked to stress resilience.

Leaf regions in barley exhibited spatial variation in heat stress responses, likely due to differences in developmental processes such as cell division and maturation across zones like the base, middle, and tip. Comparing expressed genes between leaf base and leaf tip under control conditions, we identified 622 genes specific to the leaf base and 731 genes specific to the leaf tip. These region-specific genes were enriched for distinct biological functions, where base-specific genes were primarily associated with transport and localization while tip-specific genes were largely linked to environmental stress responses in both control and heat conditions (Supplementary Figure 7 and 8). To compensate for increased investment in defense under heat stress, plants can coordinate spatially distinct transcriptional programs within a single leaf, integrating processes such as growth regulation and nutrient transport with localized stress response mechanisms. For example, *HORVU.MOREX.r3.1HG0086090* was moderately downregulated in the leaf base under heat stress (mean CPM = 303.9, log2 fold change = -0.924, adjusted p-value = 4.28e^−8^; Supplementary Figure 9). Given that the *Arabidopsis* ortholog PHO2 of *HORVU.MOREX.r3.1HG0086090* transports phosphate from root to shoot to support growth, this suggested heat-induced suppression of developmental processes. In addition, we also found that among 40 genes with significant interaction effects between leaf regions and temperature, two transporter genes – *HORVU.MOREX.r3.5HG0428260*, an ortholog of SWEET17 (sugar transporter), and *HORVU.MOREX.r3.7HG0703490*, an ortholog of LHT1 (amino acid transporter) in *Arabidopsis* – exhibited spatially temperature-responsive patterns. These results suggested that spatial coordination of growth regulation and metabolite transport contributes to the adaptive response to heat stress in cereal crops like barley. Combined with phenotypic evidence of region-specific photoprotective traits such as NPQ kinetics, our study highlights a potential growth-defense trade-off modulated by developmental status and underscores the significance of spatial context in understanding how plants allocate resources under thermal stress.

## Materials and Methods

### Experimental Design and Plant Growth Conditions

A total of 160 (including Morex as a check genotype) barley accessions from the International Barley Core Collection (BCC) (Muñoz-Amatriaín *et al*., 2014) were sown in PRO-MIX® BX medium amended with Multicote® 14-14-16 fertilizer and Micromax® micronutrients. Seeds were planted in 1.5-inch diameter, 8.25-inch-deep cone-tainers arranged in 98-cell trays (7 rows × 14 columns) at the Jack Dalrymple Agricultural Research Complex, North Dakota State University (46°53′29.70″ N, 96°48′0.91″ W). For every two columns, one Morex plant was planted to serve as the check line. Each full experiment in control or heat condition consisted of two trays and was conducted five times. To control potential sequential effects during phenotyping, planting arrangements varied across experiments. For instance, genotype entry 1 was placed first in replicate 1, whereas genotypes 30, 60, 90, and 120 were used as the first position in later replicates of experiments. Additionally, rack positions in the greenhouse were rearranged daily to reduce spatial environmental effects within the greenhouse before moving them to the growth chamber. Each rack was placed in an individual FLL small flow tray (Stuewe & Sons, Inc.) to ensure even and adequate daily watering. A 16-h photoperiod (06:00-22:00) was maintained during plant growth, and supplemental lighting was provided by five 600W High Pressure Sodium lights and four 680W LED lights in a checkerboard pattern in the greenhouse environment. At 10 days after planting, plants on two trays representing one experimental replicate were transferred to a Conviron GEN2000 growth chamber for a 3-day temperature treatment. To maintain soil moisture, each tray was bottom-watered with a shallow layer of water, ensuring only the base of each cone was submerged. Heat and control conditions were maintained at 33°C/26°C and 23°C/16°C (day/night) respectively under a 16/8-hour light/dark photoperiod and 50% relative humidity for 72 hours. Phenotype data were directly collected in the daytime after approximately 72 hours of treatment in growth chamber environments.

### SPAD measurements

To measure chlorophyll contents, the second fully extended leaf from each plant was collected and the relative amount of chlorophyll content in the base, middle, and tip sections of each leaf (Figure 1a) was recorded using a SPAD-502 Plus chlorophyll meter (Spectrum Technologies Inc., Aurora, IL).

### Chlorophyll fluorescence imaging data collections

In each experimental replicate, approximately 14-18 plants were transferred as a batch to a completely dark room for at least 15 minutes of dark adaptation following SPAD measurements. This dark adaptation was for resetting the photosynthetic apparatus to its baseline state, ensuring accurate chlorophyll fluorescence imaging data collection by eliminating the effects of prior light exposure. From each batch, the same fully extended second leaf was detached from the plant and taped abaxial down onto black paper, ensuring the abaxial surface was in contact with the paper, alongside a reference genotype Morex to control for batch effects and ensure consistency across samples collected at different time points. Images of the leaves taped onto black paper were captured using the top-view camera of the PlantExplorer Pro+ system (PhenoVation B.V., Netherlands) in the dark room. Each batch of leaves in the imaging chamber underwent an extra one minute of dark adaptation where both the minimum (F_o_) and maximum (F_m_) chlorophyll fluorescence values were extracted, followed by 5 minutes of illumination at approximately 650 µmol m^−2^ s^−1^ using a 3,000K white light source with a peak wavelength (λ_max_) of 630 nm. A 5.4 minutes of dark treatment was immediately initiated following the light exposure, incorporating saturating flashes of 3,200 μmol m^−2^ s^−1^. During the light period, maximum fluorescence measured under illuminated conditions (F_m_′) measurements were taken at intervals of 30 seconds, specifically at 30, 60, 90, 120, 150, 180, 210, 240, 270, and 300 seconds. Immediately after completing the light measurements, the dark treatment commenced. Measurements of F_m_′ during the dark-recovery treatment were recorded at 65 seconds, followed by intervals of 125, 185, 245 and 305 seconds.

### Phenotype value extractions from chlorophyll fluorescence images

To understand the spatial phenotypic variation per single leaf, five identical ROIs in the dimensions of 60 pixels x 60 pixels were evenly placed from tip to base of each individual leaf. Mean phenotypic values of chlorophyll fluorescence traits including AriIdx, ChlIdx, NDVI, F_o_, F_m_, and F_m_’ were calculated within each ROI using the software CropReporter^TM^ (v.5.8.7-64b, PhenoVation, Netherlands).

### Generating NPQ kinetics and derived traits

The NPQ values for each ROI per leaf were calculated using the formula: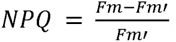, where F_m_ represents the maximal chlorophyll fluorescence recorded after a saturating pulse in dark-adapted leaves, while F_m′_ denotes the maximal fluorescence measured during actinic light exposure. The NPQ curve of each ROI was generated by connecting sequential NPQ data points and was manually inspected to remove 77 outliers including 47 in heat and 30 in control. NPQ induction and relaxation kinetics were quantified by fitting fluorescence data to previously published models (Sahay *et al*., 2024), resulting in the extraction of 14 NPQ kinetic traits from this chlorophyll fluorescence imaging dataset.

### Phenotype data processing

The BLUE values per genotype per trait was obtained using *lme4* R package v.1.1.35.3 (Bates *et al*., 2015) with the following formula,

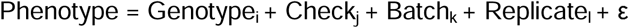

where genotype effects were modeled as fixed without an intercept, allowing separate means to be estimated for each genotype. Check line effects were salso modeled as fixed, while batch and replicate were treated as random effects. For each trait, the broad-sense heritability was calculated by treating the genotype effect as a random factor, representing the proportion of total variance attributed to genetic differences among genotypes. Broad-sense heritabilities were estimated using the following equation: 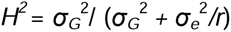, where σ*_G_*^2^ is the total genotype variance, σ*_e_*^2^ is the residual variance and *r* is the mean number of replicates per genotype within each treatment and ROI (Eltaher *et al*., 2021; Sahay *et al*., 2024).

### Genotype Data

Raw SNP data included 2,417 individual lines and their genomic positions were retrieved from T3/Barley (https://triticeaetoolbox.org/barley/; (Morales *et al*., 2022). SNPs with minor allele frequencies (MAF) < 5% or missing data rate over 90% of individuals were removed, retaining 6,574 SNPs for analysis. Hapmap genotype data was converted to variant call format (VCF) using TASSEL 5.0 (Bradbury *et al*., 2007). Missing data per genotype was imputed using Beagle (beagle.06Aug24.a91.jar; (Browning *et al*., 2018).

### Resampling Model Inclusion Probability based GWAS

Fixed and random model Circulating Probability Unification (FarmCPU) model of GWAS was implemented using rMVP v.1.0.8 (Yin *et al*., 2021; Liu *et al*., 2016) to identify association between each individual trait and SNP. To control covariate effects in the association analysis, population structure and kinship were estimated using the MVP.Data.PC and MVP.Data.Kin functions in the rMVP package, respectively. For each trait, the RMIP approach was implemented by randomly selecting 90% of total accessions for running GWAS for 200 times (Valdar *et al*., 2009). Of each individual GWAS running, the p-value threshold of 7.6e^−6^ (0.05 / the number of SNPs) was applied for multiple testing corrections using the Bonferroni method. For each SNP, RMIP was defined as the proportion of 200 GWAS runs in which the SNP was identified as significant. SNPs with an arbitrary threshold of RMIP ≥ 0.500 were considered significant. SNP and gene Browser Extensible Data (BED) files were then sorted by chromosome and position, then processed with bedtools closest -d (Quinlan and Hall, 2010) to identify the nearest gene and its distance to each significant SNP. Gene annotations were retrieved from *Ensembl Plants* release 60 (Howe *et al*., 2020) using the *Hordeum vulgare* Morex V3 pseudomolecule genome assembly. For genes without descriptions from the initial query, additional annotations were obtained from GrainGenes (Yao *et al*., 2022), using descriptions from the mRNA feature attributes. Orthologs of barley genes relative to *Arabidopsis* genes were retrieved from *Ensembl Plants* using the database’s built-in predicted orthologs in *Arabidopsis thaliana* (TAIR 10) (Howe *et al*., 2020).

### RNA-seq data generation

All seedlings of the ‘Morex’ genotype were grown in the growth chamber under the same control conditions described in the phenotyping experimental design. Fourteen days after planting, seedlings assigned to the heat treatment were exposed to a 4-hour daytime period at 33 °C, while control plants remained under standard growth conditions. For both treatments, fully expanded second leaves were collected from two to three plants per biological replicate and pooled, with three biological replicates in total. From these pooled leaves, the tip, middle, and base regions were dissected separately, flash-frozen in liquid nitrogen and stored at -80°C. Total RNA was extracted from each sample using the RNeasy Plant Mini Kit (Qiagen, USA) following the manufacturer’s recommendation. RNA concentration was measured using a NanoDrop One/OneC UV-Vis spectrophotometer (Thermo Fisher Scientific Inc., USA). Quality control was assessed through multiple criteria, including RNA quantification, agarose gel electrophoresis, RNA Quality Number (RQN) analysis, 260/280 and 260/230 absorbance ratios. Non-stranded RNA-Seq libraries were prepared and sequenced on the Illumina NovaSeq X Plus platform using a 2×150 bp paired-end configuration.

### RNA-seq data processing

Raw RNA-seq data were preprocessed to remove remaining adapters and low-quality reads using fastp v0.24.0 (Chen, 2023). The reference genome for barley Morex version 3 was obtained from Phytozome v13, while gene annotations were retrieved from *Ensembl Plants* release 60. The genome index was constructed using HISAT2 v2.2.1 (Kim *et al*., 2019) and preprocessed RNA-seq reads were aligned to the indexed reference genome using the same tool. The alignment Sequence Alignment Map (SAM) file was converted to Binary Alignment Map (BAM) format, sorted, and indexed using SAMtools v.1.21 (Li *et al*., 2009). Raw read counts per transcript were then calculated using HTseq v.2.0.5 (Li *et al*., 2009; Putri *et al*., 2022) with parameters of “-s no, -t exon, -i Parent”. The longest transcript isoform was selected to represent each gene model. The normalized expression value (referred to here as CPM) per gene was calculated using the fpm() function in DESeq2 after estimating size factors with the estimateSizeFactors() function. Genes with mean CPM > 1 across all samples were considered expressed. Principal component analysis was performed on log_₂_(CPM + 1) transformed expression values across expressed genes to visualize sample clustering patterns using R function prcomp().

Expressed genes were used to identify DEGs and genes with significant interaction effects using DESeq2 (Love *et al*., 2014) based on three modeling strategies: i) condition-specific responses (heat vs. control) were assessed independently within each leaf region using the model design ∼ Condition, with the control condition as the reference; ii) region-specific genes (base, middle and tip) were identified within each condition using the model design ∼ Region, specifying either the middle (middle vs. base, middle vs. tip) or base region (base vs. tip) as the reference. In both analyses in i) and ii), DEGs were defined as genes with adjusted p-values < 0.05 and absolute log_₂_ fold change ≥ 1; iii) genes with significant interaction effects between region and temperature were identified using the model: design ∼ Condition + Region + Condition:Region and genes with adjusted p-values < 0.001 were considered as significant.

### K-means clustering

A union of 1,833 DEGs across the three leaf regions was scaled per gene to a range of [-1, 1] using min– max normalization of log_2_ fold changes. The normalized profiles were then used for k-means clustering to identify the optimal number of expression clusters. To determine the optimal number of clusters (k), we computed the within-cluster sum of squares (WSS) for k values ranging from 1 to 12 using the kmeans() function in R. We then calculated the ratio of WSS at k to WSS at k-1 to assess the relative gain in compactness with each additional cluster. The optimal number of clusters was defined as the largest k-1 value before a noticeable drop-off in the rate of WSS reduction (Supplementary Figure 10). Based on this approach, the optimal number of clusters was determined to be 8.

### GO term enrichments

Curated GO annotations corresponding to the Morex v3 gene models were obtained from CyVerse (Wimalanathan *et al*., 2018), and all expressed genes from this study were used as the background set. GO enrichment analysis was performed using GOATOOLS v1.4.12 (Klopfenstein *et al*., 2018) for specific studied gene groups and GO terms with Bonferroni-corrected p-values ≤ 0.05 were considered as significantly enriched.

## Author contributions

Z.K.L. conceived and designed the experiments. E.C.F., Z.K.L. S.Y. and Z.H.L collected the data, E.C.F., G.T. W.D. and Z.K.L. analyzed the data, and E.C.F., and Z.K.L. wrote the manuscript. All authors read and approved the final manuscript.

## Acknowledgements

This work was supported by the Agricultural Research Service Grant #58-3060-3-016 from the U.S. Department of Agriculture, American Malting Barley Association, Montana Wheat & Barley Council and the North Dakota State University Start-up fund to Z.K.L. The findings and conclusions in this preliminary publication have not been formally disseminated by the U. S. Department of Agriculture and should not be construed to represent any agency determination or policy. This work used resources of the Center for Computationally Assisted Science and Technology (CCAST) at North Dakota State University, which were made possible in part by National Science Foundation Major Research Instrumentation (MRI) Award No. 2019077. The authors thank Rong Zhou for thoughtful discussions about photoprotection mechanisms under abiotic stress and Zachary Springer for support in soil preparation for this study.

## Conflict of Interest Statement

The authors declare no competing interests.

## Data Availability Statement

RNA-seq data from three leaf regions under control and heat conditions in barley genotype Morex, each with three biological replicates, have been deposited in the NCBI Sequence Read Archive (SRA) under BioProject ID PRJNA1275958.

## Notes

### Competing Interest Statement

The authors have declared no competing interest.

